# Chemical imaging unveils mitochondria as the major site of medicinal biguanide accumulation within cells

**DOI:** 10.1101/2024.01.02.573981

**Authors:** Lei Wang, Xianrong Zeng, Yanjie Li, Wanyu Hao, Zijing Yu, Luxia Yao, Yongdeng Zhang, Zhaobin Wang, Lianfeng Wu

## Abstract

Metformin (MET), a commonly prescribed medication for managing type 2 diabetes, has demonstrated various beneficial effects beyond its primary anti-diabetic efficacy. However, the mechanism underlying MET activity and its distribution within organelles remain largely unknown. In this study, we integrate multiple technologies, including chemical labeling, immunostaining, and high-resolution microscopy imaging, to visualize the accumulation of MET in organelles of cultured cells. To achieve this objective, an alkynylated MET probe is developed that preserves biological activity similar to biguanide drugs. As determined by biorthogonal chemical labeling and imaging, the MET probe selectively localizes to substructures within cells, contrasting with its probe control. Furthermore, the MET probe can be competitively and efficiently washed out through biguanide administration, demonstrating the specific activity of this probe in monitoring the cellular dynamics of biguanide drugs. Our results indicate that the MET probe can reach near-saturated concentrations within two hours and is rapidly eliminated within an additional two hours once the exogenous source of the drug is removed. Furthermore, we reveal that the MET probe primarily accumulates in mitochondria, particularly within the mitochondrial matrix, and has a minor presence in other organelles, such as lysosomes and endosomes. Together, this study provides the first view of the MET subcellular localization and lays the foundation for future investigations on its molecular targets and mechanisms of action in promoting human health.

## Introduction

In addition to its primary antidiabetic usage, metformin (MET) has shown potential for the management of other prevalent chronic diseases, such as aging, obesity, cardiovascular disease, neurodegenerative diseases, and cancer (He, 2020; Rena et al., 2017; Soukas et al., 2019). Despite its widespread use in the clinic and extensive study in basic research, the mechanisms underlying how and where metformin exerts its effects remain poorly understood. Previous pharmacokinetic investigations have reported that oral metformin is absorbed into the bloodstream from the gut, primarily acts in the liver and is predominantly excreted through the kidneys (Graham et al., 2011; Rena et al., 2013). Recent isotope imaging studies using C^11^-MET demonstrated that MET is also taken up and accumulates in other tissues, including heart, salivary glands, skeletal muscle, and adipose tissue (Breining et al., 2018; Gormsen et al., 2016; Iversen et al., 2017); therefore, MET may be universally distributed in cells of various tissues. However, the locations of MET at the intracellular level remain unclear.

Mitochondria, as a powerhouse of eukaryotic cells, have been widely associated with the development and progression of various chronic metabolic diseases (Bhatti et al., 2017). Notably, numerous studies have suggested that MET directly or indirectly acts through mitochondrial inhibition (Bridges et al., 2023; Guo et al., 2017; Tramonti et al., 2021; Wu et al., 2016). Multiple mitochondrial proteins have been reported as direct targets of MET, such as mitochondrial complex I, mitochondrial glycerol phosphate dehydrogenase (mGPD) (Madiraju et al., 2014) and CYP3A4 (Guo et al., 2017), strongly indicating that mitochondria are target organelles of metformin. Additionally, direct targets of MET have been proposed in other organelles, including PEN2 (Ma et al., 2022) from lysosomes and HMGB1 from the nucleus (Yuan et al., 2020). However, the location in which MET targets and accumulates within cells remains elusive.

Through recent advancements in chemical labeling, immunostaining, and high-resolution microscopy imaging, the dynamics of a compound within subcellular structures can be comprehensively analyzed. For instance, the Cu-catalyzed alkyne-azide cycloaddition (CuAAC) reaction is a recently developed biorthogonal reaction that covalently connects compounds to biological molecules within cells (Liang and Astruc, 2011). This reaction has been widely used for in situ fluorescence imaging of compound dynamics in cells at subcellular resolution and target fishing from cell lysates (Pang et al., 2022). However, this analysis has never been used to demonstrate the dynamic distribution of the MET.

In this study, we developed a MET probe based on the principle of the CuAAC reaction, which has similar biological effects as biguanide drugs on the activation of AMPK signaling and the inhibition of mTORC1 activity. This probe was linked with an Alexa Fluor 647 dye in the CuAAC reaction, enabling specific visualization of MET molecules compared to its backbone compound control in HeLa cells. The probe was competitively and markedly washed out of the cells when exogenous MET or phenformin was administered together. A rapid accumulation and excretion of MET was observed within two hours. Through chemical labeling and immunostaining approaches, we revealed that MET was predominantly enriched in mitochondria and had a minor presence in lysosomes and endosomes among various organelles. This explicit map of MET presence within subcellular structures will help researchers reveal its mechanisms of action and identify its direct targets.

## Results

### A MET probe bearing similar and specific activity to biguanide drugs

To visualize the intracellular distribution of MET within cells, we developed and synthesized the MET probe by introducing an alkyne group in the para-position of the benzene ring of phenformin (Figures 1A and S1A-1G). Through utilizing the backbone of phenformin, the active center of the biguanidine moiety was not interrupted. A backbone compound including the phenyl group but not the biguanidine moiety of phenformin was also synthesized as a control of the MET probe (Figure 1A). First, using the widely accepted signaling readouts downstream of MET response pathways, we investigated whether the MET probe maintained activity similar to biguanide drugs. These MET response outcomes included the activation of the AMP-activated protein kinase (AMPK) pathway (Ma et al., 2022) and inhibition of the target of the rapamycin complex 1 (mTORC1) pathway (Wu et al., 2016), which are indicated by the phosphorylation levels of acetyl-CoA carboxylase (pACC) and phosphorylated ribosomal protein (pS6), respectively (Figure S2A). The results demonstrated that our MET probe could stimulate those readouts in a dose-dependent manner (Figure S2A). Remarkably, the MET probe exhibited similar activity in activating AMPK or inhibiting mTORC1 signaling to phenformin when used at the same concentrations (both 200 and 500 μM and MET at concentrations of 1 and 10 mM) (Figure S2B). Collectively, these results demonstrated that the MET probe maintained similar activity to biguanide drugs and almost the same as the effect of phenformin.

**Figure 1.**
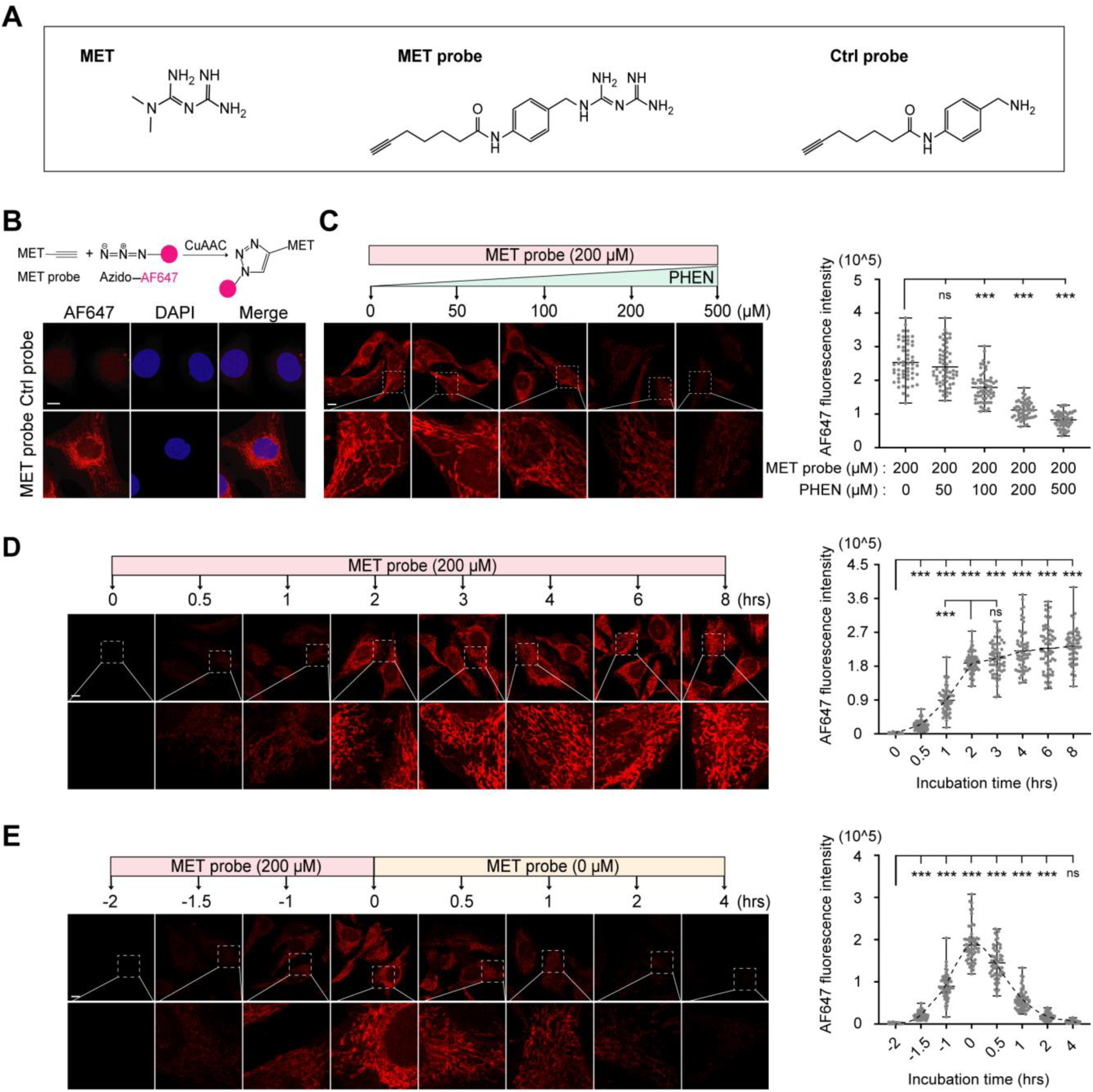
A MET probe bearing similar activity as biguanide drugs. (A) The chemical structure of metformin (MET), the MET probe, and the control (Ctrl) probe. (B) Representative confocal images of the intracellular MET or Ctrl probe. MET or Ctrl probes were covalently linked to the fluorescent dye AF647 via the CuAAC reaction (n = 3 replicates). DAPI, nuclear staining. The upper graphic depicts the principle of the CuAAC reaction. Scale bars, 10 μm. (C) Representative confocal images of the intracellular MET probe upon competition with different doses of phenformin (PHEN) (upper) for 2 hours and the corresponding quantification of AF647 intensity (right). Images in the lower panel show an enlarged view of the dashed rectangular area in the upper images. Each data point (n = 54-60 from 24 images per group) was normalized to the group without PHEN treatment. Multiple unpaired t tests were applied. Scale bar, 10 μm. (D) Representative confocal images of the intracellular MET probe (left) and the corresponding quantification of AF647 intensity (right). Images in the lower layer show an enlarged view of the dashed rectangular area. Each data point (n = 60 from 24 images per group) was normalized to the indicated 0 time point. Scale bar, 10 μm. (E) Representative confocal images of the intracellular MET probe (upper) under the indicated incubation strategy above the images and the corresponding quantification of AF647 intensity (right). Images in the lower layer show an enlarged view of the dashed rectangular area. Each data point (n = 60 from 24 images per group) was normalized to the indicated 0 time point. The gray dashed line connects the means of each data point. Multiple unpaired t tests. Scale bar, 10 μm. For all statistical data (C, D and E), multiple unpaired t test analysis was applied. Each dot represents a calculated fluorescence intensity value within the selected regions. Data are presented as the means ± SEMs. No significance, ns; *p<0.001*, ***. CuAAC, copper(I)-catalyzed azide-alkyne cycloaddition.

Through the CuAAC biorthogonal reaction, the synthesized probes could be covalently bound with the substrate dye azido-Alexa Fluor 647 (AF647) in cultured cells (Figure 1B). Via this reaction, the distribution of those probes within cells could be visualized (Figure 1B). Strikingly, we observed that the MET probe specifically accumulated in certain subcellular structures, including mitochondria and nuclei, while the control probe was primarily found in the nucleus. To validate whether the MET probe could specifically mimic the performance of biguanides, we conducted competition assays using phenformin and metformin. Remarkably, we found that the MET probe was competitively washed off at 200 μM by supplementation with phenformin and metformin; intensity reduced by approximately 50% and 75% by 200 µM and 500 µM phenformin, respectively, and an approximately 50% decreased in intensity was attained by 10 mM metformin (Figure 1C and Figure S3). Collectively, these findings suggest that the MET probe exhibits similar and specific properties to biguanide drugs.

### Dynamic and subcellular distribution of biguanides within cells

Next, we investigated the dynamics of the MET probe in HeLa cells and found that the probe efficiently accumulated in cells and reached saturated levels in two hours (Figure 1D). This timing for the probe to reach the peak concentration was surprisingly similar to that of metformin orally administered in patients with T2D (Sun et al., 2011), further suggesting the specificity of the probe for representing biguanide drugs in illustrating their dynamics. Furthermore, we observed that the probe exited the cells rapidly and was reduced to a nearly undetectable level 2 hours after its exogenous supply was removed (Figure 1E). These results indicated a fast turnover rate for the influx and efflux of biguanide drugs at the cellular level.

To determine the location of the biguanide drugs at the organelle level, we performed colocalization analysis using the MET probe and a set of antibodies that specifically mark different organelles (Figure 2). The results showed that biguanides exhibited an evident and major accumulation in mitochondria and a minor presence in other organelles, such as lysosomes, peroxisomes, endoplasmic reticulum, and endosomes. These results suggest that biguanides may act through multiple targets localized in distinct organelles, particularly targets in mitochondria, to exert pleiotropic effects.

**Figure 2.**
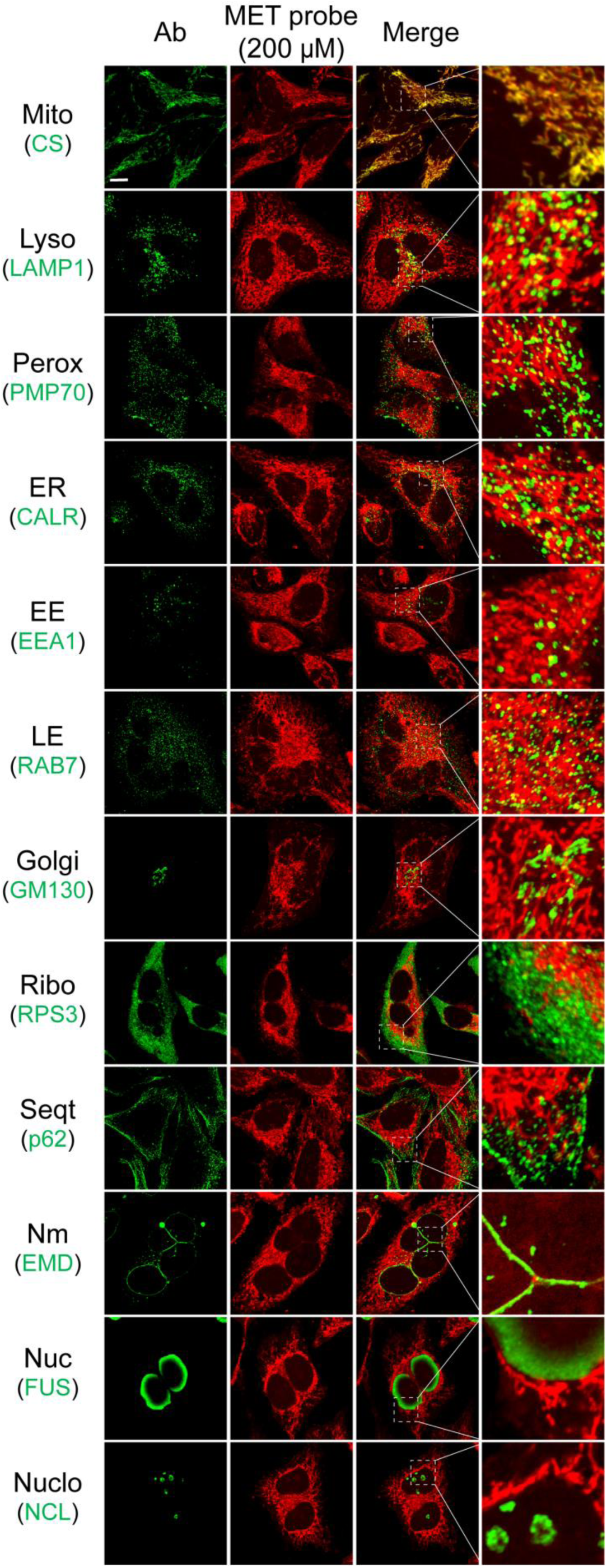
Organelle distribution of the medicinal biguanide. Representative images of the MET probe (red) and the organelles (green) marked with the indicated antibodies (Ab) in HeLa cells (n = 2). The right panel shows the enlarged region of the dashed box from the corresponding images on the left. Mitochondria (Mito), Lysosome (Lyso), Peroxisome (Perox), Endoplasmic reticulum (ER), Early endosome (EE), Late endosome (LE), Golgi apparatus (Golgi), Ribosome (Ribo), Sequestome (Seqt), Nuclear membrane (Nm), Nuclear matrix (Nuc) and Nucleolus (NCL). Scale bar, 10 μm.

### Biguanide primarily accumulates in the mitochondrial matrix

To further explore the specific localization of the MET probe within mitochondria, we performed super-resolution imaging analysis using different markers of mitochondrial substructures. These markers included MitoSOX Red, a fluorescent dye staining the mitochondrial matrix, and antibodies targeting the matrix protein citrate synthase (CS), the inner mitochondrial membrane protein ATP synthase F1 subunit alpha (ATP5A1), and the outer mitochondrial membrane protein translocase of outer mitochondrial membrane 20 (TOM20) (Figure 3). The images depicted that the signal of the MET probe perfectly aligned with the boundary of the MitoSOX Red signal and completely covered the signal regions of CS and ATP5A1. In contrast, the MET probe signal was completely contained by the signal boundary of TOM20. These results demonstrate that biguanides primarily accumulate in the mitochondrial matrix, strongly indicating that the direct targets of biguanide drugs in mitochondria are most likely in the mitochondrial matrix.

**Figure 3.**
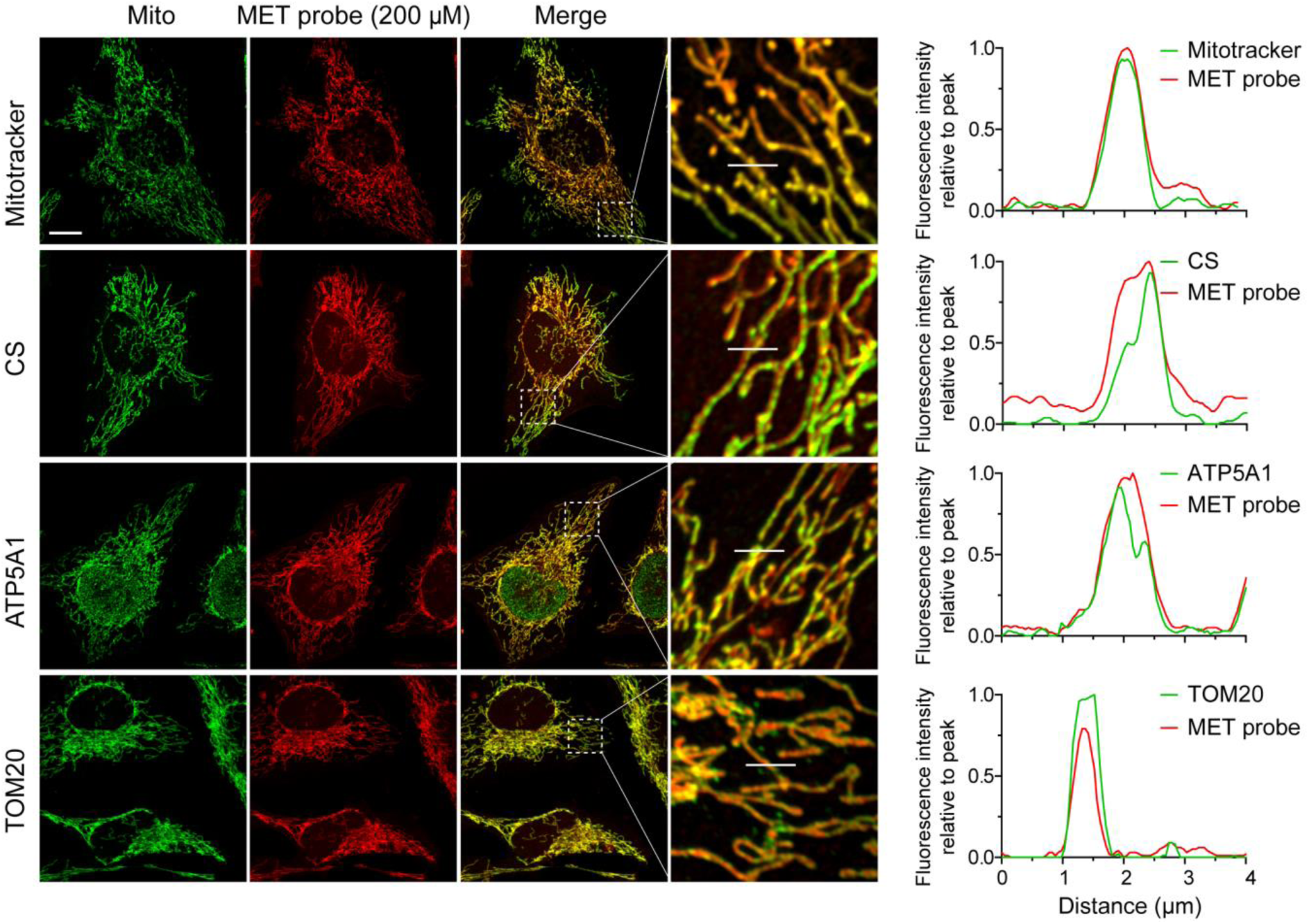
Mitochondrial matrix as the major reservoir of biguanide accumulation in cells. Representative images of the MET probe within mitochondria of HeLa cells co-stained with MET-AF647 (red) and the indicated fluorescent-conjugated antibodies (green, n = 2). MitoTracker dye, CS and ATP5A1 and TOM20 mark the mitochondrial interstitium-matrix, matrix, inner and outer membranes, respectively. The fluorescence intensity analysis of green and red channels in each column along the x-axis corresponding to the white line in the enlarged images (right). Scale bar, 10 μm.

## Discussion

The old saying that “Seeing is believing” always leads to the most compelling illustrations for the nature of a biological event. The mystery about what organelles the panacea drug metformin approaches or targets within cells remains elusive, although the drug has been prescribed worldwide and suggested to have direct targets within mitochondria and lysosomes (Bridges et al., 2023; Ma et al., 2022; Viollet et al., 2012). In the current study, we developed a chemical probe that did not interrupt the biguanidine group of biguanide drugs and was experimentally demonstrated to exhibit comparable activity to biguanide drugs. Using this MET probe, the influx and efflux of biguanide at a relative therapeutic concentration (200 µM) (Frid et al., 2010) within cells could be analyzed via imaging approaches. The combination of chemical labeling for MET probe tracing and immunofluorescence staining for organelle marking yielded the first visualization of MET distributions at the organelle level, especially for its enrichment in mitochondria. Our findings depict the dynamics of efflux and influx within cells and the first map of MET distribution in organelles of cells, which will aid future studies on MET target search and mechanism exploration.

Our study revealed a nearly saturated mode for the entry of the MET probe into cells and its rapid excretion out of cells, both within two hours. These results may explain why cells or animals can survive through very high concentrations of MET treatment (Wu et al., 2016). Its rapid excretion property may also provide explanations for the clinical dosing strategy in which patients with T2D take MET orally two to three times a day for an effective treatment (Davidson and Peters, 1997; Lily and Godwin, 2009). This excretion mode of MET implies that metformin may utilize an immediate or transient mode of action on its targets. The competitive examinations demonstrated that the MET probe may exhibit more similar dynamic properties to phenformin, the uptake of which into cells has been reported to be more efficient and less dependent on specific transporters than metformin (Yuan et al., 2013). However, the mechanisms underlying the efflux dynamics of biguanide drugs need further exploration.

Our study also clearly showed that the medicinal biguanide is predominantly enriched in mitochondria. This corresponds with the previous finding in which its concentration reached up to 1000-fold higher within the mitochondria than the level in the extracellular medium (Owen et al., 2000). However, it remains unclear whether MET entry into mitochondria is dependent on specific transporter proteins, such as OCTs (Kimura et al., 2005; Shikata et al., 2007; Shu et al., 2007; Wang et al., 2002), or the electrochemical potential (ΔΨ) energy formed across membranes (Zorova et al., 2018). MET carrying two positive charges by its nature very likely alters the charge distribution and pH environment within the mitochondrial matrix. It would be meaningful to determine whether this biguanide drug alters mitochondrial metabolism by modulating mitochondrial charge properties, as such conditions greatly influence the activities of mitochondrial enzymes (Kitada and Ito, 2001). Our study pinpoints that MET is predominantly enriched in mitochondria despite its possible but minor presence in other vesicle trafficking vesicles, including lysosomes and endosomes, at the imaging level. This finding further underscores the significance of thoroughly investigating the mitochondrial activity of MET. Such research will ultimately unveil the mechanisms underlying MET action.

## Methods

### Chemical synthesis, validation and purification of designed probes

Unless otherwise noted, all reagents were purchased from TCI, Energy Chemical, Bide, and Leyan. All reactions were performed under atmospheric atmosphere. Solvents were purchased from J&K or Energy Chemical and used directly. The synthesized probes were both validated using nuclear magnetic resonance (NMR). NMR spectra were recorded on a Bruker spectrometer with a Prodigy broadband cryoprobe (500 MHz and 600 MHz); chemical shifts (δ) are reported in ppm downfield from tertramethylsilane, using the solvent resonance as the internal standard. High-resolution mass spectrometric analysis was performed on an ultra-performance liquid chromatography-time-of-flight mass spectrometer (Synapt-G2-Si, Waters, USA) with electron spray ionization (ESI) resources.

The Ctrl probe (aminoalkyne substrates, **S4**) preparation:

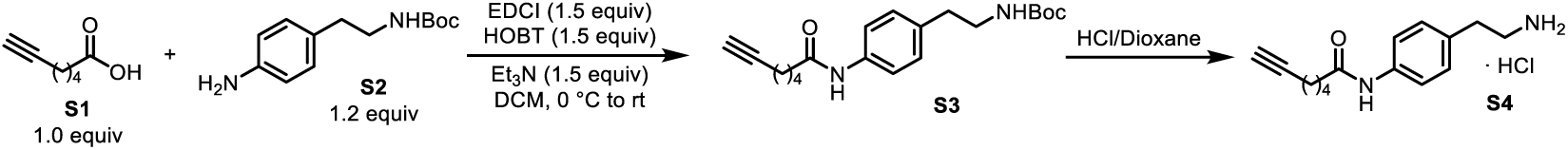

Condensation: A solution of **S1** (1.8 mL, 14.26 mmol) in CH_2_Cl_2_ (20 mL) was added to a solution of EDC·HCl (4.06 g, 21.18 mmol) and HOBt (2.88 g, 21.18 mmol) in CH_2_Cl_2_ (40 mL), followed by the addition of a solution of amines (4.0 g, 16.92 mmol) in CH_2_Cl_2_ (20 mL). The mixture was cooled to 0 °C, and Et_3_N (3.0 mL, 21.18 mmol) was added dropwise. After being slowly warmed to room temperature and stirred overnight, the reaction mixture was diluted with CH_2_Cl_2_, washed successively with water, 2N HCl, saturated NaHCO_3_, and brine, and dried over Na_2_SO_4_. Evaporation of the solvent followed by column chromatography on silica gel with EtOAc/PE (1:2) as eluent afforded **S3**: 4.81 g, 98% yield, yellow solid.

Deprotection: In an inert gas atmosphere, **S3** (4.81 g, 13.96 mmol) was placed in a 100 mL round bottom flask, and HCl/dioxane (4 M, 50 mL) was added at 0 °C. The reaction was stirred overnight at room temperature. The ice bath was removed, and the solution was stirred overnight at room temperature. The solvent was removed under reduced pressure, and the product was filtered off and washed with diethyl ether to produce **S4** (3.91 g, 99% yield, white solide). **S4** was used as the Ctrl probe for the indicated experiments or subsequent transformations without further purification.

**S4**: 3.91 g, 99% yield, white solid.

^1^H NMR (500 MHz, DMSO) δ 10.00 (s, 1H), 8.08 (s, 2H), 7.56 (d, *J* = 8.4 Hz, 2H), 7.16 (d, *J* = 8.4 Hz, 2H), 2.97 (m, 2H), 2.83 (m, 2H), 2.76 (t, *J* = 2.6 Hz, 1H), 2.31 (t, *J* = 7.4 Hz, 2H), 2.18 (td, *J* = 7.1, 2.6 Hz, 2H), 1.66 (m, 2H), 1.47 (m, 2H).

^13^C NMR (126 MHz, DMSO) δ 170.9, 138.0, 131.7, 128.8, 119.3, 84.3, 71.3, 40.0, 35.7, 32.4, 27.5, 24.3, 17.5.

HRMS (ESI) m/z [M + H]^+^ calcd for C_15_H_21_N_2_O: 245.1654, found: 245.1651.

MET probe (Phenylethylbiguanide dihydrochloride, **P**) preparation:

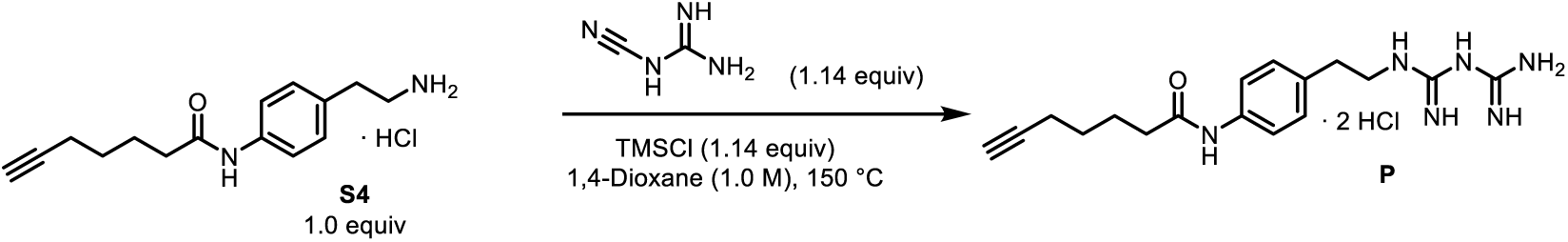

In an atmospheric gas atmosphere, **S4** (200 mg, 0.7 mmol) was placed in a 20 mL sealed tube, and dicyandiamide (67 mg, 0.8 mmol), TMSCl (101 μL, 0.8 mmol) and solvent 1,4-dioxane (0.7 mL) were added successively and stirred overnight at 150 °C. After being slowly warmed to room temperature, the reaction mixture was diluted with EtOAc, the product was filtered out, washed with EtOAc and CH_2_Cl_2_ to prepare the product, and purified using liquid phase preparation chromatography.

**P**: 180 mg, 64% yield, 72% purity, white solid.

^1^H NMR (500 MHz, MeOD, 55 °C) δ 7.48 (m, 2H), 7.20 (m, 2H), 3.44 (m, 2H), 2.83 (m, 2H), 2.38 (t, *J* = 7.5 Hz, 2H), 2.23 (m, 3H), 1.81 (m, 2H), 1.59 (m, 2H).

^13^C NMR (126 MHz, MeOD, 55 °C) δ 174.3, 161.5, 160.5, 138.2, 135.9, 130.1, 121.8, 84.7, 69.6, 43.9, 37.3, 35.9, 29.2, 25.9, 18.8.

HRMS (ESI) m/z [M + H]^+^ calcd for C_17_H_25_N_6_O: 329.2090, found: 329.2092.

MET probe purification: Semipreparative liquid chromatography (Waters Pre150, C18 reversed-phase HPLC column) was used to purify the MET probe. RP-HPLC analyses were performed over an XBridge® Prep C18 5 µm OBD^TM^ column (19 × 150 mm) using Pre150 Semi-Preparative Liquid Chromatography (Waters). Solvent A was ddH_2_O. Solvent B is a mixture of methanol (Sinopharm, 40064292) and acetonitrile (Fisher Chemical, A998) at a 1:1 ratio. Solvents A and B are ultrasonically treated to remove bubbles before use. The sample concentration of the crude MET probe, which was dissolved in methanol, was approximately 10 mg/mL, and 100 μL of solution was loaded into the chromatographic column in each injection. After four consecutive injections, the pipeline system was re-equilibrated. The retention time of the MET probe was near 6 min, and the maximum absorbance of the MET probe was at a wavelength of 238 nm. The target fraction was collected by the automatic collection system within the set time range. After the solvent was removed by spin drying, the powder of the MET probe obtained had a purity of 96% according to mass-spectrum identification. Purified drug powder was dissolved in DMSO and stored at −20 ℃ prior to use.

### Cell culture

HeLa cells were regularly cultured in RPMI-1640 medium supplied with 10% fetal bovine serum, 100 U/mL penicillin, and 0.1 mg/mL streptomycin and were grown at 37 °C with 5% CO_2_. Five percent fetal bovine serum was used for Western blot analysis. All cells were checked for *mycoplasma* contamination before use.

### Western blot

After drug treatment, the cells were lysed in hot (70 °C) SDS lysis buffer (10 mM Tris-HCl, 2 mM EDTA, 1% SDS). Protein quantification of tested samples was determined using a BCA kit. Protein samples were boiled in 1 x SDS loading buffer prior to SDS‒PAGE. Twenty micrograms of protein per lane was loaded into a precast PAGE gel for electrophoresis separation in MOPS running buffer. Protein bands were transferred onto 0.22 m PVDF membranes for immunoblotting. Membranes were blocked in 5% BSA or 5% skim milk in TBST (TBS buffer containing 0.1% tween-20) buffer for 2 hours at RT and then incubated with primary antibody overnight at 4 °C or RT for 3 hours. HRP-conjugated secondary antibodies were incubated with the membranes for 1 hour at RT, and the signals were detected with ECL reagent by a chemiluminescence imaging system. Fiji software was used to analyze the gray values of the bands on the WB membrane.

### Chemical labeling and imaging

After incubation for 16 hours, HeLa cells were seeded into a 24-well plate at a density of 0.25 x 10^5 cells per well with cover a slide. The cell surface was gently washed with a pipette to remove the loosened attached or dead cells, the cover slide was transferred to a new well containing fresh treatment medium, and then the cells were incubated with a control or MET probe accordingly. After incubation, the cells were fixed with 4% PFA for 30 min at RT and then immediately quenched with 100 mM freshly prepared glycine solution in 1 x PBS buffer. The quenched cells were permeabilized with 0.1% Triton X-100 prepared in PBS buffer for 30 min at RT and washed three times with PBS with gentle rotation (5 min each time). The following procedures were performed in a dark area. During washing, 1 mL CuAAC reaction solution (25 μM AF647, 1 mM CuSO4, 2 mM BTTP and 100 mM ascorbic sodium) was prepared. The cells were immersed in 300 µl of reaction solution and placed on a horizontal shaker with 50 rpm rotation for 1 hour at RT. After the reaction, the cells were washed five times with PBS buffer (10 min each time), stained with DAPI and mounted in H-1000 mounting medium. The slides were stored at 4 °C for observation within 2 days or at −20 °C for longer preservation. A Zeiss LSM 900 or Olympus FV3000 Laser Confocal system was used to acquire images, and the images were processed by using instrument configured software, Fiji software (win64), or an Imaris workstation. High-resolution images were taken using an FV3000. Superresolution z-stack images (3-5 μm thickness) were captured under the AiryScan module of Zeiss LSM900. Colocalization analysis and calculation of the Pearson correlation coefficient (PCC) and Mander’s overlap coefficient (MOC) were performed using Imaris software. The signal border projected from MitoTracker, CS, ATP5A1, and TOM20 was measured by Fiji software (convert RGB color or 8 bit> LUT-edges), and the fluorescence intensity of MET-AF647 or secondary antibody-conjugated fluorescent dye was counted by Fiji software (convert RGB color or 8 bit> adjust Color Threshold > Analyze, Measure).

To attain colocalization imaging between the MET probe and MitoTracker, MitoTracker Red CMXRos (stock solution is 250 x, dissolved in DMSO) was diluted into a medium and incubated with cells for 30 minutes before fixation. The following processing steps are the same as described above. Colocalization imaging between the MET probe and protein markers of the oganelles. All procedures were performed in a dark area. The antibody staining procedures followed the washing steps after the CuAAC reaction. MET-AF647-marked cells were incubated in a blocking solution (5% goat serum, 0.3% Triton X-100 in PBS at pH=8.0) for 1 hour at RT with horizontal rotation (50 RPM), followed by incubation with the primary antibody overnight at 4 °C or at RT for 3 hours. Cells were washed three times (5 min each) in PBS and then incubated with secondary antibodies for 1 hour at RT. After the cells were washed three times with PBS, they were mounted, preserved, imaged, or analyzed as described above.

### Quantification of MET-AF647 fluorescence intensity

For fluorescence intensity quantification, Fiji software was used. Each image with a full size of 1024 pixels x 1024 pixels was first converted into an 8-bit grayscale format, adjusting the threshold level at the value of 5 for the lower threshold level and 255 for the upper threshold level. Two to four regions with a size of 150 pixal x 150 pixal (approximately 50% background and 50% signal region) in each given image were randomly chosen for integrated density (IntDen) statistics of the MET-AF647 fluorescence. Eight confocal images from one biological replicate were used for fluorescence quantification. Each experiment was conducted independently at least three times.

### Statistical analysis

Statistical significance was analyzed with multiple unpaired t tests. Only comparisons with *p < 0.05* were defined as significant. Each imaging dynamics experiment was repeated in three independent experiments. The WB experiment was repeated three independent times, and the band intensity was quantified by Fiji software. The chemical structure was drawn by ChemOffice-Suite-2019. Statistical analyses were performed using GraphPad Prism.

## Author contributions

L.F.W. and L.W. conceived the idea, designed the experiments, and coordinated activities among authors; L.W., X.R.Z., Z.J.Y., Y.J.L., W.Y. H., and L.X.Y. conducted the experiments; L.F.W., L.W., and Z.B.W. designed the structure of the probes; X.R.Z. completed the synthesis and characterization of the MET probe under supervision of Z.B.W.; Y.D.Z. and Z.J.Y. performed and analyzed the super-resolution imaging; L.F.W. and L.W. analyzed the data and wrote the manuscript with inputs from all others.

## Acknowledgments

We thank the “Instrumentation and Service Center for Molecular Sciences” and “Biomedical Research Core Facilities” at Westlake University for their technical support. We also thank Dr. Yinjuan Chen and Ke Wang (research assistants) for their help in chemical identification and purification. This study was supported by the National Natural Science Foundation of China (32071151 and 32271357 to L.F.W; the Natural Science Foundation of Zhejiang Province (2022XHSJJ005 to L.F.W.), the Postdoctoral fellowship (2022M722842 to L.W.), and the institutional funding (L.F.W and Z.B.W).

## Resource availability

Reasonable requests for further information and resources as well as reagents should be directed to and will be fulfilled by Dr. Lianfeng Wu.

## Data and code availability

Source data for the graphs are provided in Data S1. No original code was generated in this study.

---

**Figure S1.**
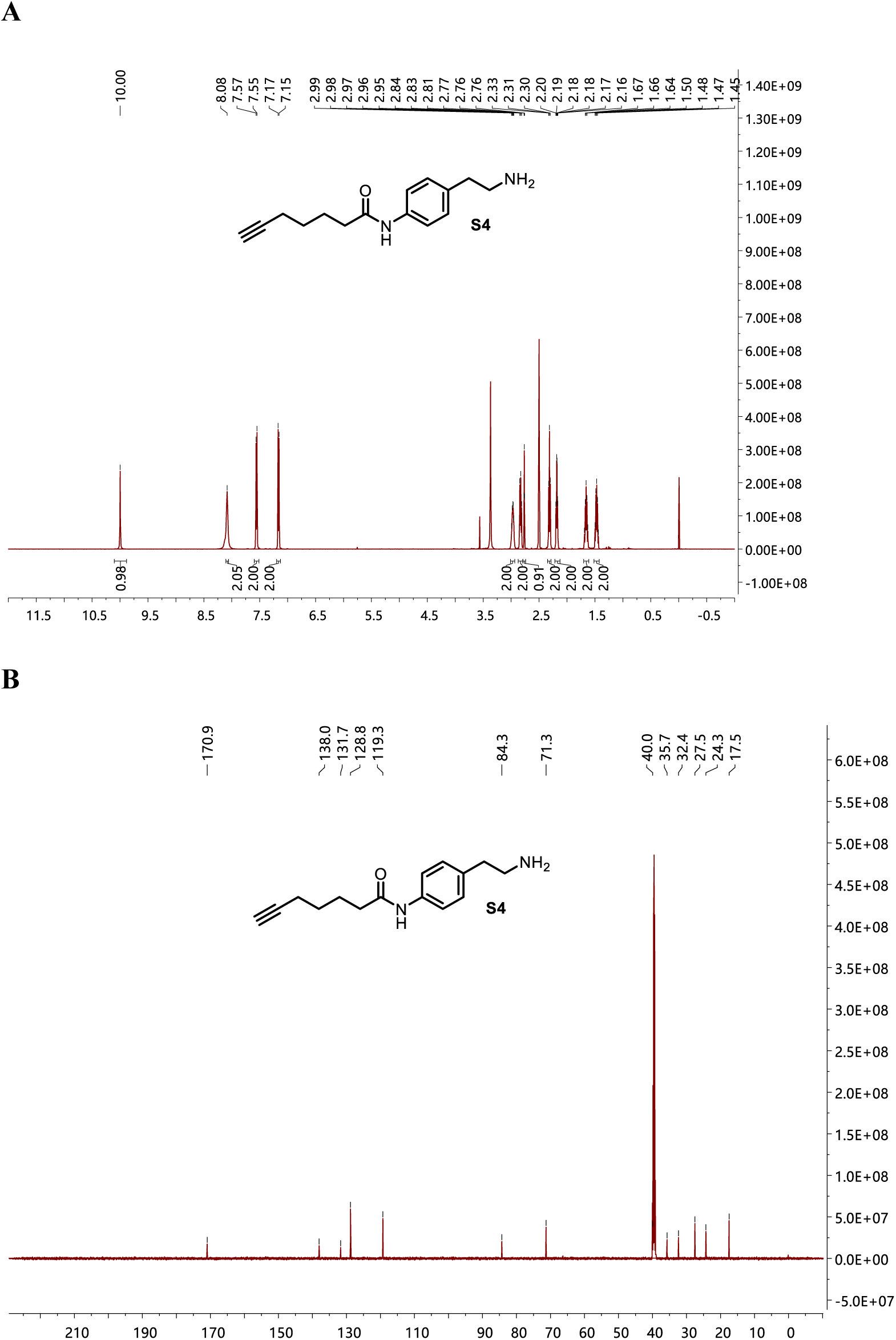
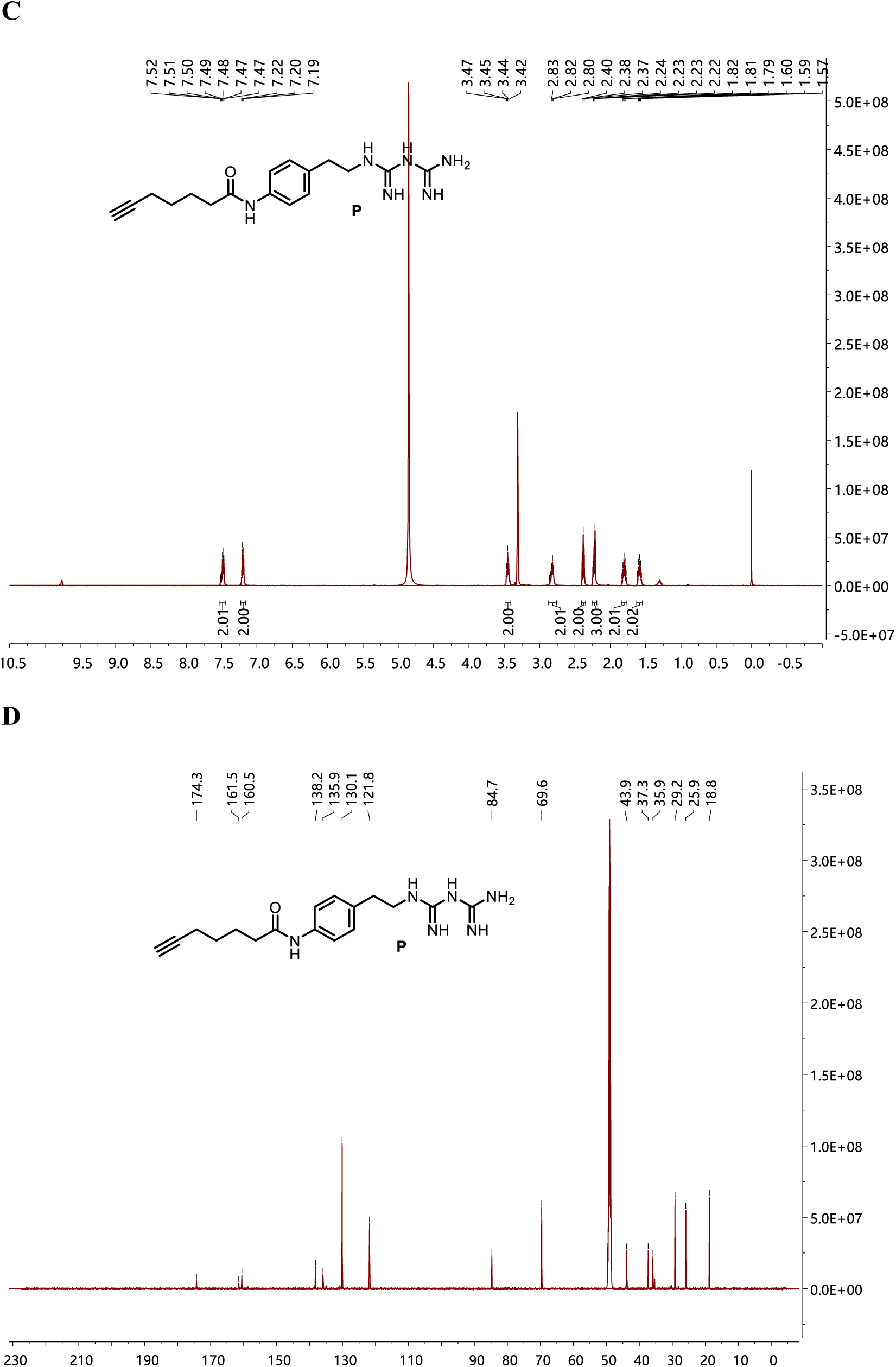
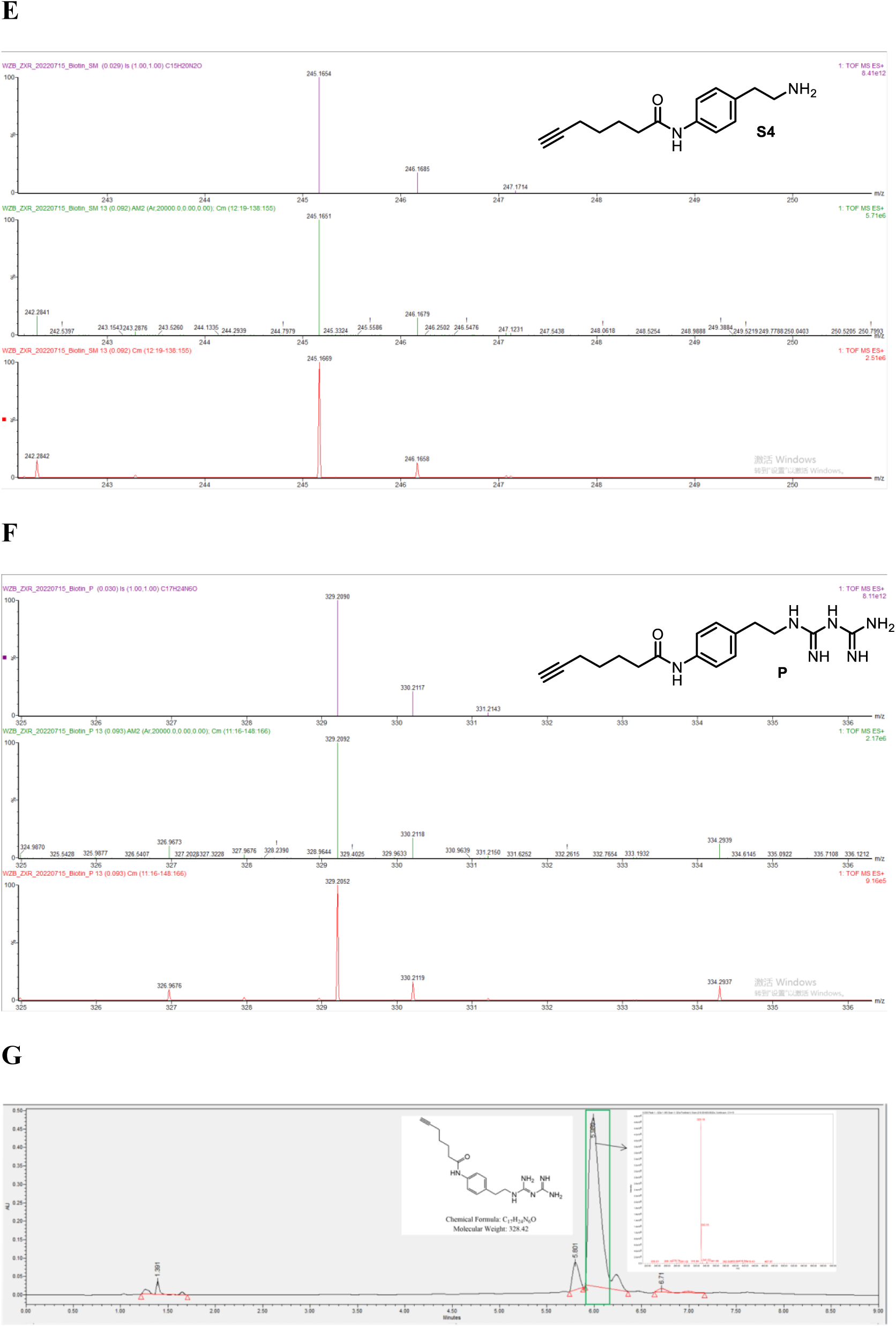
Validation and purification of the synthesized Ctrl and MET probes. (A-D) The ^1^H NMR (500 MHz, DMSO) and ^13^C NMR (126 MHz, DMSO) spectra of the Ctrl (A, B) and MET (C, D) probes. (E, F) High-resolution mass spectrometry (HRMS) results for validation of the Ctrl (E) and MET (F) probes. (G) Chromatogram results for the purification of the MET probe.

**Figure S2.**
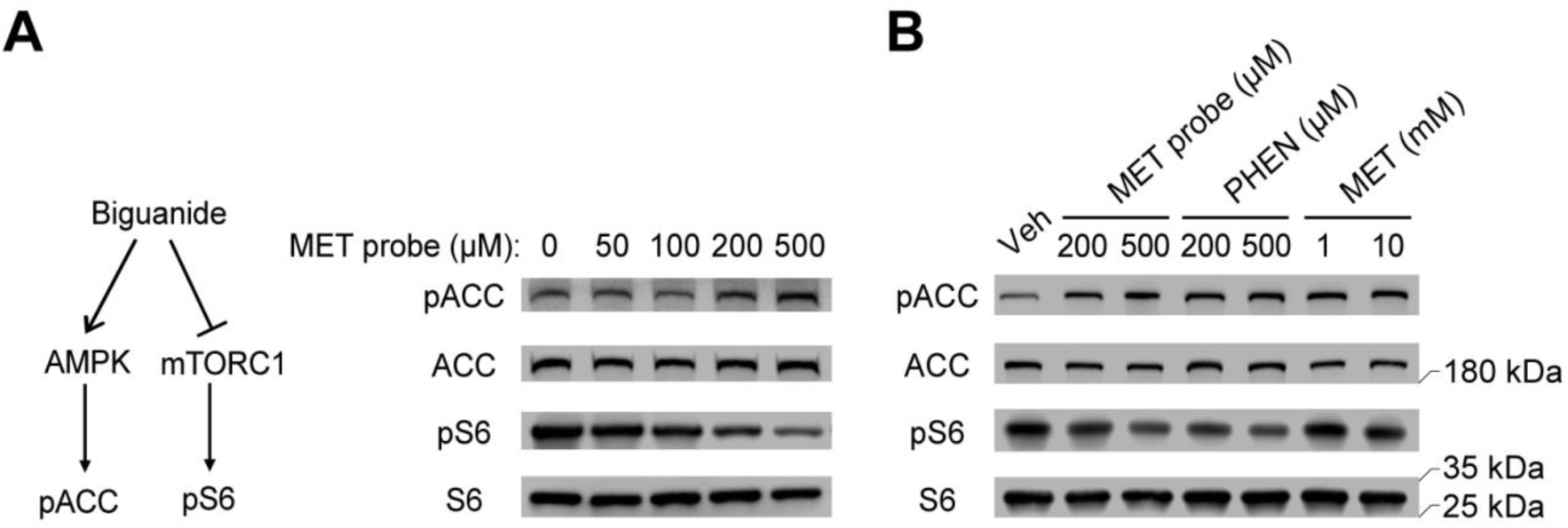
The MET probe exerts similar effects as biguanide drugs on the activation of AMPK and the inhibition of mTORC1 signaling. Representative Western blot results are shown for the effects of the MET probe on inactivated AMPK activation and mTORC1 inhibition in HeLa cells after an 8-hour treatment (n = 3). The diagram (left) presents the widely recognized effects of biguanide drugs on AMPK activation and mTORC1 inhibition. The levels of phosphorylated ACC (pACC) and phosphorylated S6 (pS6) were used to indicate changes in activity of AMPK and mTORC1, respectively. Representative Western blot data displaying the effects of the MET probe, phenformin (PHEN) and metformin (MET) on AMPK activation and mTORC1 inhibition at different concentrations in HeLa cells (n = 3). The compound treatment was conducted for 8 hours.

**Figure S3.**
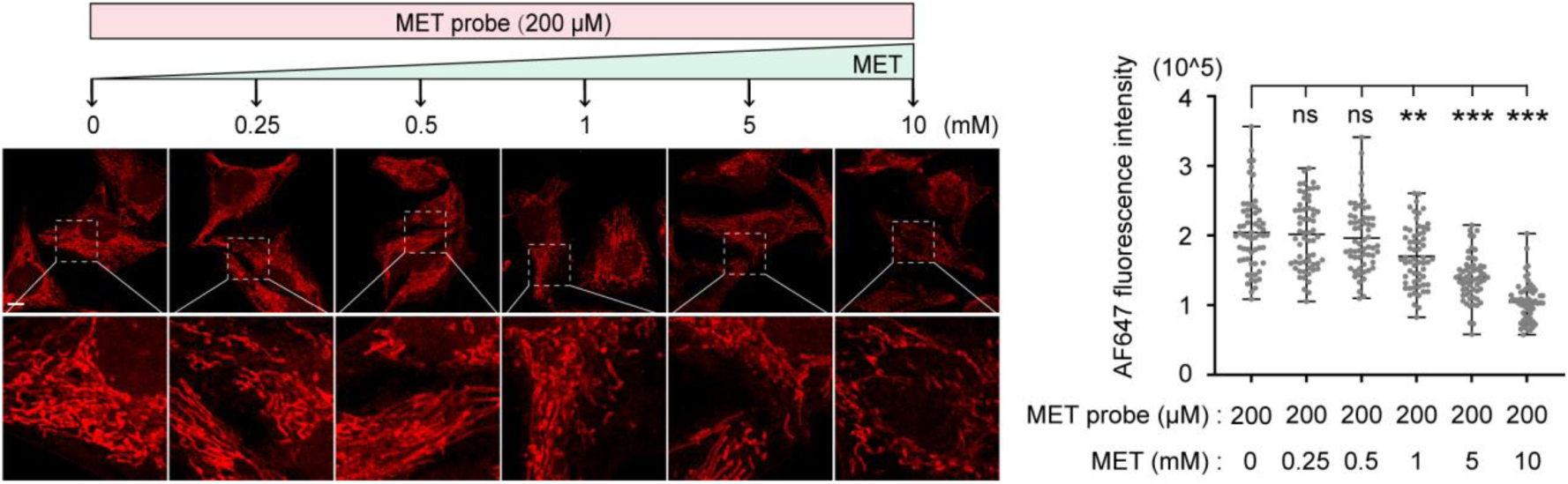
The MET probe can be competitively washed out of the cells by metformin administration. Representative images of the intracellular MET probe upon competition with different doses of MET for 2 hours (left) and the corresponding quantification of AF647 intensity (right). Images in the lower layer show an enlarged view of the dashed rectangular area of the upper images. Each data point (n = 60 from 24 images per group) was normalized to the indicated 0 time point. Scale bar, 10 μm. For all statistical data, multiple unpaired t test analysis was applied. Each dot represents a calculated fluorescence intensity value within the selected regions. Data are presented as the means ±SEMs. No significance, ns; *p<0.01*, **; *p<0.001*, ***.

